# Caldendrin represses neurite regeneration via a sex-dependent mechanism in sensory neurons

**DOI:** 10.1101/2021.07.26.453831

**Authors:** Josue A. Lopez, Annamarie Yamamoto, Joseph T. Vecchi, Jussara Hagen, Amy Lee

## Abstract

Caldendrin is a calmodulin-like Ca^2+^ binding protein that is expressed primarily in neurons and regulates multiple effectors including Ca_v_1 L-type Ca^2+^ channels. Here, we tested the hypothesis that caldendrin regulates Ca_v_1-dependent pathways that repress neurite growth in dorsal root ganglion neurons (DRGNs). By immunofluorescence, caldendrin was localized in medium- and large-diameter DRGNs. Consistent with an inhibitory effect of caldendrin on neurite growth, neurite initiation and growth was enhanced in dissociated DRGNs from caldendrin knockout (KO) mice compared to those from wild type (WT) mice. In an *in vitro* axotomy assay, caldendrin KO DRGNs grew longer neurites via a mechanism that was more sensitive to inhibitors of transcription as compared to WT DRGNs. Strong depolarization, which normally represses neurite growth through activation of Ca_v_1 channels, had no effect on neurite growth in DRGN cultures from female caldendrin KO mice. Remarkably, DRGNs from caldendrin KO males were no different from those of WT males in terms of depolarization-dependent neurite growth repression. We conclude that caldendrin opposes neurite regeneration and growth, and this involves coupling of Ca_v_1 channels to growth-inhibitory pathways in DRGNs of females but not males. Our findings suggest that caldendrin KO mice represent an ideal model in which to interrogate the transcriptional pathways controlling neurite regeneration and how these pathways may differ in males and females.

## INTRODUCTION

Dorsal root ganglion neurons (DRGNs) are pseudo-unipolar neurons with a peripherally directed neurite that senses information about modalities, such as touch and temperature, which transmits this information into the spinal cord via a centrally directed neurite. In contrast to neurons whose cell bodies reside in the central nervous system, DRGNs retain the ability to regenerate their peripheral neurite following injury (Scheib & Hoke, 2013). However, the slow rate of regrowth and incomplete re-innervation of target tissues remain significant barriers for therapies to restore sensory function in individuals suffering from peripheral nerve damage (Huebner & Strittmatter, 2009). Despite insights into the molecular mechanisms underlying the functional diversity of DRGNs (Mecklenburg et al., 2020; Wu et al., 2021; Zheng et al., 2019), comparatively little is known about the factors that control the regenerative growth of DRGN neurites.

In many neurons, Ca^2+^ ions are essential second messengers that can serve as either positive or negative regulators of neurite growth. In cortical and hippocampal neurons of immature rodents, Ca^2+^ influx through voltage-gated Ca_v_1 (L-type) channels promotes the growth and arborization of dendrites via mechanisms that involve calmodulin (CaM)-dependent protein kinase and transcription (i.e., “excitation-transcription” coupling). In the context of DRGNs, however, Ca_v_1 channels trigger pathways that repress neurite regeneration in DRGNs: Ca_v_1 antagonists facilitate DRGN neurite growth *in vitro* (Enes et al., 2010; Huebner et al., 2019) and genetic disruption of Ca_v_1 expression promotes recovery from peripheral nerve injury *in vivo* (Enes et al., 2010). Within DRGNs, the mechanisms whereby Ca_v_1 channels repress neurite growth may involve the transcription of neurite growth inhibiting genes (Enes et al., 2010) or alterations in membrane trafficking and cytoskeletal proteins in the advancing tip of the neurite (*i*.*e*., growth cone) (Gomez & Zheng, 2006; Henley & Poo, 2004).

Caldendrin is a member of a family of Ca^2+^ binding proteins (CaBPs) that are related to calmodulin (CaM) and highly expressed in neuronal cell-types of the brain, retina, and inner ear (Haeseleer et al., 2004; Laube et al., 2002; Tippens & Lee, 2007; Yang et al., 2016). In cochlear spiral ganglion neurons, caldendrin regulates Ca_v_1 channels and their coupling to the repression of neurite growth by strong depolarization (T. Yang et al., 2018). To probe the involvement of caldendrin in Ca_v_1-dependent mechanisms controlling neurite regeneration, we characterized the expression of caldendrin in DRGNs and analyzed neurite growth in cultured DRGNs from wild-type (WT) and caldendrin knockout (KO) mice. We find that caldendrin is prominently localized in medium- and large-diameter DRGNs and represses neurite regeneration via a transcription-dependent mechanism that involves Ca_v_1 channels in females but not in males. Our results reveal an unexpected role for caldendrin in controlling sexually dimorphic pathways converging on neurite growth in sensory neurons.

## METHODS

### Animals

All experiments were performed in accordance with guidelines set by the Institutional Animal Care and Use Committee at the University of Iowa. The mice (4-8 week old males and females) were housed under a standard 12-h light/dark cycle with access to food and water *ad libitum*. The generation and characterization of caldendrin KO mice (RRID: MGI: 5780462) was described previously (Kim et al., 2014) and were maintained on a C57BL/6 (Envigo) background.

### Immunofluorescence of DRGs

Lumbar DRGs (L4-L6) were harvested from male and female adult WT C57BL/6 mice and immersed in a 4% paraformaldehyde solution (PFA) made in phosphate-buffered saline (PBS, Cat #AAJ19943K2, ThermoFisher) and stored at 4 °C overnight prior to transfer to 15% sucrose solution in PBS for an additional 24 h. The fixed tissue was molded in optimal cutting temperature compound (Sakura Finetek) and 15-μm thick sections were obtained using a cryostat (Leica). Sections were mounted on charged slides and dried on a slide warmer for 5 min. Sections were then subject to rinsing, blocking, and antibody incubations at room temperature (RT). After rinsing in PBS and incubating in blocking buffer (10% normal goat serum, Cat #PCN5000 and 0.2% Triton™ X-100 Cat #9002-93-1, ThermoFisher, in PBS) for 30 mins, the sections were incubated for 1 h in the following primary antibodies diluted 1:1000 in blocking buffer: chicken polyclonal antibody anti-NF200 (RRID: AB_2313552, Aves); chicken polyclonal antibody anti-Peripherin (RRID:AB_10785694, Aves Labs); rabbit polyclonal antibody against caldendrin (UW72) (Haeseleer et al., 2000). Sections were washed 3 times with PBS for 10 min followed by incubation with secondary antibodies diluted 1:500 in blocking buffer: goat anti-Chicken Alexa Fluor®546 (RRID: AB_2534097, ThermoFisher) and Goat anti-Rabbit Alexa Fluor®488 (RRID:AB_2534096, ThermoFisher). Sections were then washed 3 times with PBS for 10 min each and mounted using Fluoromount-G® (SouthernBiotech) and allowed to cure for 24 h in the dark before imaging. DRGs from at least 4 mice were processed in 4 independent experiments. At least 3 sections were analyzed using an Olympus Fluoview 1000 confocal laser scanning microscope with a 20 x lens and FluoView software (RRID: SCR_014215). The numbers of cells exhibiting colocalized caldendrin and NF200 or peripherin fluorescence was determined by researchers blinded to the labeling conditions and genotype.

### Western Blots

Five lumbar WT and caldendrin KO DRGs were harvested from 4-8-week-old male and female mice, flash frozen in liquid nitrogen, and lysed in lysis buffer (50 mM Tris-HCl pH 7.4, 150 mM NaCl, 0.5% Triton X100, 0.5% n-Dodecyl-beta-Maltoside detergent (Thermofisher Cat# 89902), Protease Inhibitor Cocktail (Roche cOmplete, EDTA-free (Cat# 05056489001)) and 1 mM PMSF. Plastic pestles were used to homogenize the pellet in a 1.5 ml microfuge tube. Lysates were cleared by spinning down samples at 14 x 1000 rcf at 4 °C for 10 min. NuPAGE™ LDS Sample Buffer (4X) (Cat# NP0007, ThermoFisher) was added to lysates which were then incubated at 65 °C for 10 min. The lysate (25% of total) was loaded into a 4-20% Tris-Glycine gel (Invitrogen Cat# XP04200BOX), run in Tris-Glycine SDS Running buffer (Novex Life technologies Cat# LC2675) and transferred overnight using Tris-Glycine Transfer Buffer (Novex Life technologies Cat# LC3675). Blots were incubated in blocking buffer (5% fat free milk in TBST) for 30 min prior to incubation for 1 h in primary antibodies diluted in blocking buffer: Goat anti-mouse GAPDH (RRID: AB_1078991, Sigma-Aldrich) 1:1000, goat anti-mouse GAP43 (RRID: AB_2533123, Thermo Fisher Scientific) 1:1000, rabbit polyclonal antibody against caldendrin and goat anti-mouse β-tubulin (RRID: AB_396360, BD Biosciences) 1:3000. Chemiluminescent detection was accomplished with a Li-COR Odyssey Imager by exposing the blot for 10 min and Image Studio lite was used to prepare images.

### DRGN cultures and immunofluorescence

DRGN cultures were prepared as previously (Lin & Chen, 2018). One day before culture preparation, 14 mm round glass cover slips (Deckglaser) were placed in 24 well plates (Cat# 229123, Celltreat) and coated with poly-L-ornithine (0.1 mg/ml in 10 mM borate buffer, pH 8.4, P4957, Sigma) overnight at 37°C. Coverslips were rinsed with Milli-Q H_2_O three times and coated with laminin (20 μg/ml in ultra-pure H_2_O, 11243217001, Sigma) for 1 hr at 37°C. Mice were anesthetized with isoflurane and subjected to cervical dislocation and decapitation. 20-30 DRGs were harvested into Neurobasal™ media (Cat#21103049, ThermoFisher) and DRGNs were dissociated in 0.125% Trypsin-EDTA (Cat #25200056, ThermoFisher) and 0.1% collagenase type I (1 mg/mL in HBSS without calcium or magnesium (-/-)), (Cat #17100, ThermoFisher) for 40 min. After the enzyme incubation, 10% horse serum was added to inactivate the proteases, the cell suspension was centrifuged at 500 rcf for 5 min. The neurons were resuspended in Neurobasal™ media supplemented with 2.5% L-glutamine (Cat #25030081, ThermoFisher) and 2% N-21 (Cat #SCM081, Millipore Sigma), plated at a density of 5 ganglia per well, and cultured for 18, 24 and 48 h in a humidified incubator at 37 °C with 5% CO_2_. In the replating assay (Saijilafu et al., 2013), DRGNs were cultured for 72 h prior to dissociation with TrypLE™ Express (Cat#12604013, ThermoFisher) for 1 min and replating and culture for an additional 24 h. Dichlorobenzimidazole riboside (40 µM, Cat# D1916 Sigma-Aldrich) or nocodazole (50 µM, Cat# M1404 Sigma-Aldrich) diluted in dimethylsulfoxide (DMSO) or DMSO control solution was added to the medium either before or after replating. In some experiments, 40 mM KCl (K_40_) ± isradipine (Cat#2004, Tocris) or nimodipine (each at 10 μM), Cat#0600, Tocris)

After culture for 18-48 h, media was removed and the coverslips washed with PBS. All subsequent steps were done at room temperature. DRGNs were fixed by adding 4% PFA for 10 min, washed with PBS (3 times, 5 min each), incubated in blocking buffer for 30 min, and subsequently for 1 h in chicken polyclonal antibody against NF200 (RRID: AB_2313552, Aves) diluted 1:1000 in 5% normal goat serum (Cat #PCN5000, ThermoFisher), 0.5% TritonTM X-100 (Cat #9002-93-1, Fisher Scientific), Each well was washed 3 times for 5 mins with PBS followed by addition of Goat anti-Rabbit Alexa Fluor®488 (RRID:AB_2534096, ThermoFisher) (1:1000 in blocking buffer) for 1 h in the dark. The DRGs were then incubated with DAPI 1:10,000 in PBS for 5 mins. The DRGNs were washed with PBS (3 times, 5 min each). The coverslips were mounted using Fluoromount-G® (SouthernBiotech) and cured for 24 h in the dark before viewing on an Olympus BX53 microscope with a 20x objective equipped with Olympus DP72 camera and CellSens Standard imaging software (RRID: SCR_014551).

### Quantitative analysis of neurite growth

All quantitative analysis was performed by researchers blinded to experimental conditions. In DRGN cultures, only the NF200 positive neurons with DAPI-positive nuclei were included in the analyses. The % of DRGNs with neurites was determined by dividing the total number of DRGNs with a neurite at least as long as the diameter of the cell soma with the total number of DRGNs on the coverslip and multiplying by 100. n = 300-400 neurons per coverslip, 1 coverslip per animal, at least 3 WT or caldendrin KO mice per experiment). For analyses of the longest axon, images of DRGNs were acquired using an Olympus DP72 camera and CellSens Standard imaging software (RRID: SCR_014551). The longest axon was measured using Fiji (ImageJ) plug-in NeuronJ(Popko et al., 2009).

### Statistical analysis

Statistical analysis was done with Graphpad Prism 9 (RRID: SCR_002798, GraphPad Software, San Diego, CA). Data were first tested for normality by Shapiro-Wilk normality test. If the data were normally distributed, unpaired t-test or ANOVA with Sidak multiple comparison test, with a single pool variance was performed. If data were not normally distributed, Mann-Whitney test was performed. For two-way ANOVA test, with main effects only, and Tukey’s multiple comparisons test was performed with individual variances computed for each comparison. Otherwise, post hoc tests were performed, and simple main effects were reported using adjusted p value for multiple comparisons. Data are presented as mean and SEM and p<0.05 was considered statistically significant.

## RESULTS

### Caldendrin is expressed in medium and large diameter DRGNs

The CaBP1 subfamily of CaBPs includes caldendrin and the shorter CaBP1 variants (CaBP1-S, CaBP1-L; Fig. 1A) (Haeseleer et al., 2000; Seidenbecher et al., 1998). Caldendrin is the primary CaBP1 variant expressed in the nervous system and exists as 36 kDa and 33 kDa forms (Caldendrin-L and Caldendrin-S) that may arise from alternate translational start sites (Kim et al., 2014). By Western blot, only Caldendrin-L and Caldendrin-S were detected with rabbit polyclonal antibodies capable of recognizing all three variants (Kim et al., 2014) (Fig. 1B). The 33- and 36-kDa bands likely corresponded to caldendrin since they were detected in DRG lysates from WT but not caldendrin KO mice that lack expression of all 3 CaBP1 variants (Kim et al., 2014). To gain insights into which subpopulations of DRGNs express caldendrin, we performed double-labeling with antibodies against caldendrin and neurofilament 200 heavy chain (NF200) or peripherin, markers for a marker for myelinated, medium-to large-A-β fiber DRGNs and unmyelinated C-fiber and thinly myelinated A-δ fiber DRGNs, respectively (Figs. 1C,D) (Lawson & Waddell, 1991). Quantitative analyses showed that caldendrin was present in ∼92% of NF200-positive neurons and only ∼5% of peripherin-positive neurons. Thus, caldendrin is the major CaBP1 variant expressed in DRGNs and is localized mainly in medium and large-diameter DRGNs.

**Figure 1.**
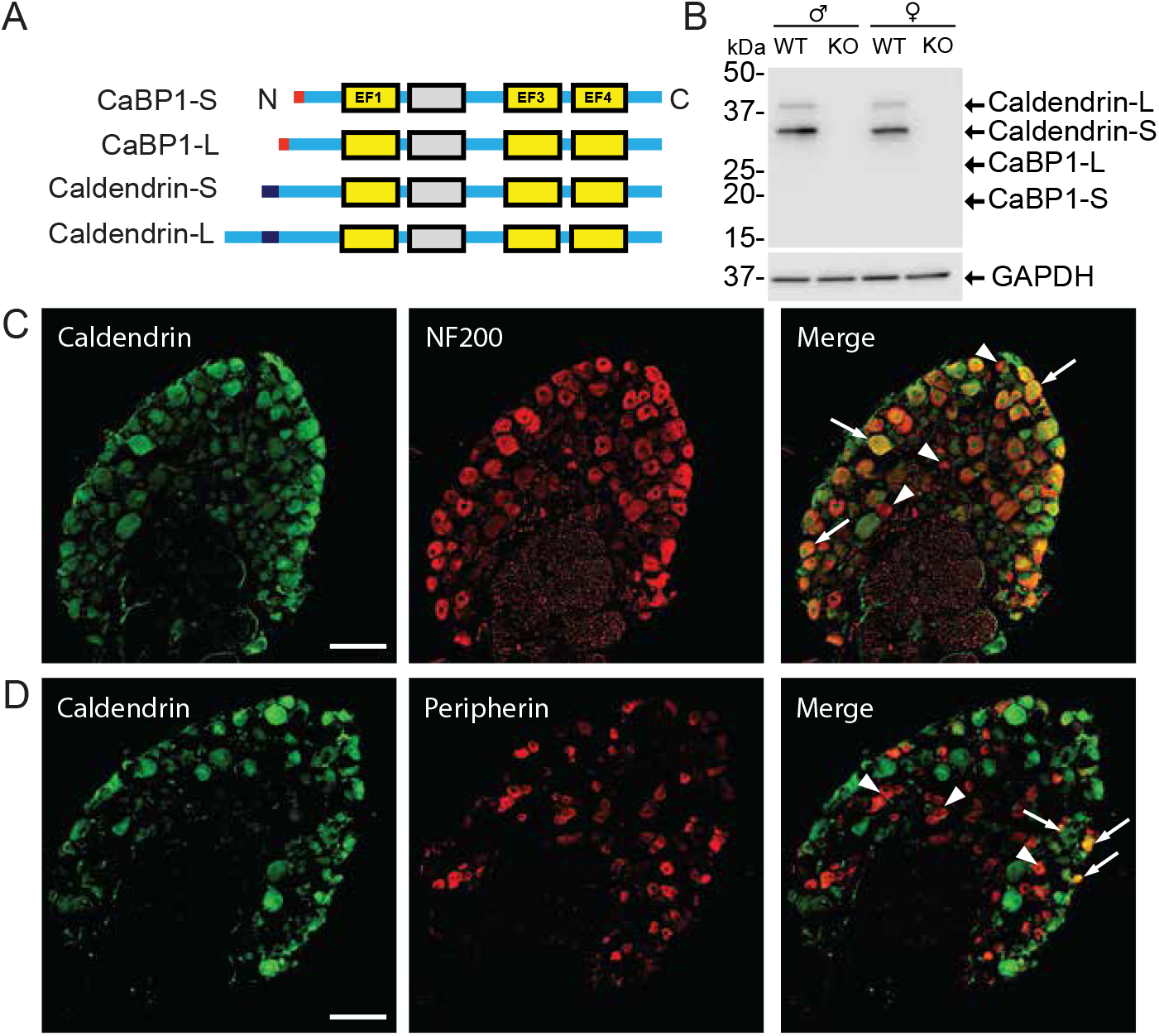
Caldendrin is the major CaBP1 variant in the DRG and is expressed primarily in myelinated DRGNs. (A) Schematic of CaBP1 splice variants containing EF-hand Ca^2+^ binding motifs and variable N-terminal regions. CaBP1-S and CaBP1-L contain an N-myristoylation site (red). Caldendrin-S and Caldendrin-L have a longer N-terminal region that includes a proline rich region (purple). (B) Western blots of DRG lysates from WT and KO mice probed with an antibody that recognizes all CaBP1 variants and ß-tubulin antibody for a protein loading control. 36- and 33-kDa bands corresponding to Caldendrin-S and Caldendrin-L were detected in WT but not in KO lysates. (C,D) Confocal micrographs from WT mouse DRG cryosections double-labeled with antibodies against caldendrin and NF200 or peripherin. In merged image, regions of colocalization are color-coded in yellow. Results are representative of 3 independent experiments. Scale bar, 100 µm.

### Caldendrin KO DRGNs exhibit enhanced neurite initiation

Based on our previous studies of neonatal cochlear spiral ganglion neurons from caldendrin KO mice (T. Yang et al., 2018), we hypothesized that caldendrin acts as a repressor of neurite growth in sensory neurons. If so, then neurite regeneration should be enhanced in DRGNs from caldendrin KO mice as compared to those from WT mice. To test this, we analyzed the percentage of neurons with neurites in cultures of DRGNs from mice. This value was significantly greater in caldendrin KO DRGNs than in WT DRGNs at 18 h and 24 h, but not at 48 h after plating in cultures from male (Fig.2B) and female mice (Fig.2B,C). There was no significant difference in the frequency distribution or median values of neurite lengths in WT and caldendrin KO cultures 24 h after plating (data not shown). These results suggest that caldendrin represses the mechanism that enables the initial but not later stages of neurite regeneration.

**Figure 2.**
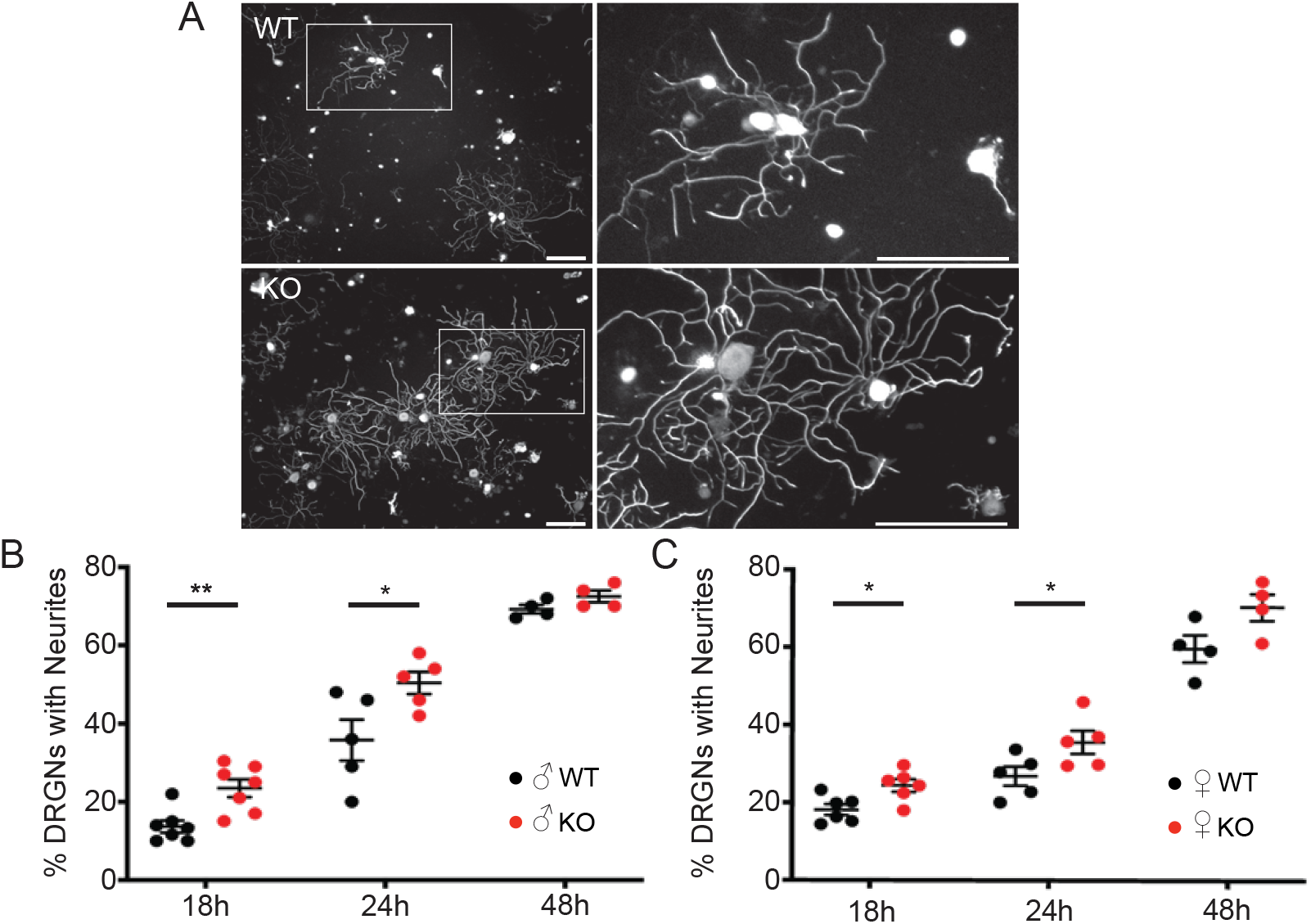
Neurite regeneration is enhanced in DRGNs of caldendrin KO than WT mice. (A) Representative images of WT and KO DRGNs immunolabeled with NF200 antibodies after 24 h in culture. Boxed region in left panels is shown at high-magnification in right panels. Scale bars, 200 μm. (B,C) Percent of DRGNs with neurites measured as described in Materials and Methods for cultures from WT and KO males (B) and females (C) processed after the indicated times in culture. Each point represents the average obtained from a single culture (n=300-400 DRGNs/culture), error bars represent SEM. **, p=0.004; *, p ≤0.05, compared to WT by unpaired t-test.

### Neurite regeneration is facilitated in caldendrin KO DRGNs following in vitro axotomy

Within 24 h of dissociation and culture, DRGNs from adult rats develop compact arborizing neurites prior to an elongating, less branched phase of neurite growth by 48 h (Smith & Skene, 1997). The transition to the elongating growth phase requires gene transcription programs that can be triggered by peripheral nerve injury (Moskowitz et al., 1993; Schreyer & Skene, 1991) such that DRGNs cultured from animals subject to *in vivo* axotomy extend few neurites that rapidly elongate within 24 h (Smith & Skene, 1997). To test whether caldendrin influences the arborizing or elongating phase of neurite growth, we used an *in vitro* axotomy assay in which DRGNs from naive mice are cultured for 3 days prior to replating. The initial 3 days following dissociation mimics the conditioning response to nerve injury that requires gene transcription, such that after replating DRGNs exhibit primarily an elongating pattern of neurite growth that depends on local assembly-driven mechanisms within growth cones (Saijilafu et al., 2013). If caldendrin represses neurite regeneration following *in vitro* axotomy, then neurites should be longer in DRGNs from caldendrin KO than from WT mice upon replating. Indeed, the frequency distribution of neurite lengths was shifted rightward, and the median length of the longest neurite was higher in cultures from caldendrin KO than WT mice, regardless of sex (Fig.3A-D).

**Figure 3.**
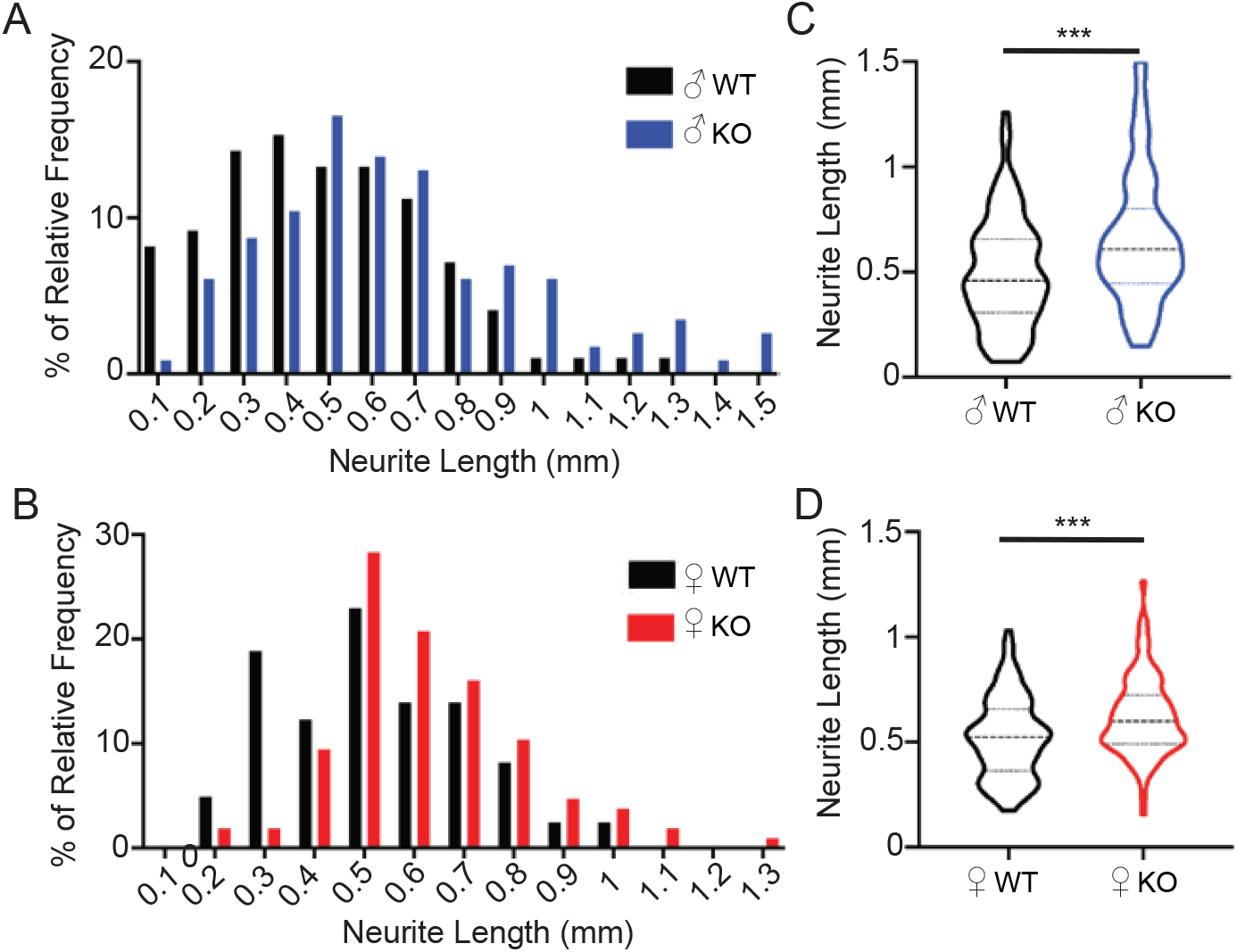
Caldendrin KO DRGNs exhibit enhanced neurite regrowth after *in vitro* axotomy. DRGNs were cultured for 3 days prior to replating and culture for 24 h prior to processing for NF200 immunofluorescence and measurement of the longest neurite per DRGN. (A,B) Frequency distributions of maximal neurite lengths in male (A) and female (B) WT and KO cultures. Data was collected from images of individual DRGNs (n = at least 32 images per culture, n = 3 independent cultures). (C,D) Violin plots of data in *A,B* with dashed and dotted lines representing the median and quartiles, respectively. ***, p=0.0002 by Mann-Whitney test.

Peripheral nerve injury induces the expression of regeneration-associated genes (RAGs) (Costigan et al., 2002), which is thought to prime DRGNs for the elongating phase of neurite outgrowth (Smith & Skene, 1997). In the replating assay, RAGs including GAP43 are upregulated in the initial 3-day culture period, similar to that which occurs following peripheral nerve injury (Schreyer & Skene, 1991). Blunting transcription with the RNA polymerase II inhibitor, 5,6-dichlorobenzimidazole riboside (DBR) before but not after replating prevents the switch from the arborizing to elongating phase of neurite growth (Fig.4A-C), whereas treatment with the microtubule destabilizer, nocodazole, administered after but not before replating suppresses cytoskeletal mechanisms involved in neurite elongation (Saijilafu et al., 2013). If caldendrin boosts transcriptional alterations that promote neurite regeneration, then DBR should be more efficacious in suppressing neurite outgrowth in caldendrin KO than WT cultures. By contrast, if caldendrin acts solely by the transcription-independent pathway, exposure of the replated DRGNs to ND should more strongly suppress neurite growth in cultures from caldendrin KO than from WT mice. We found that DBR caused a significantly greater reduction in neurite outgrowth in caldendrin KO than in WT cultures, but only when administered during the transcription-dependent phase (*i*.*e*., prior to replating) whereas nocodazole had similar effects in repressing neurite growth in the replated DRGNs from WT and caldendrin KO mice (Fig.4). These results establish that caldendrin represses neurite regeneration and growth via the transcription-dependent rather than the local-assembly dependent pathway.

**Figure 4.**
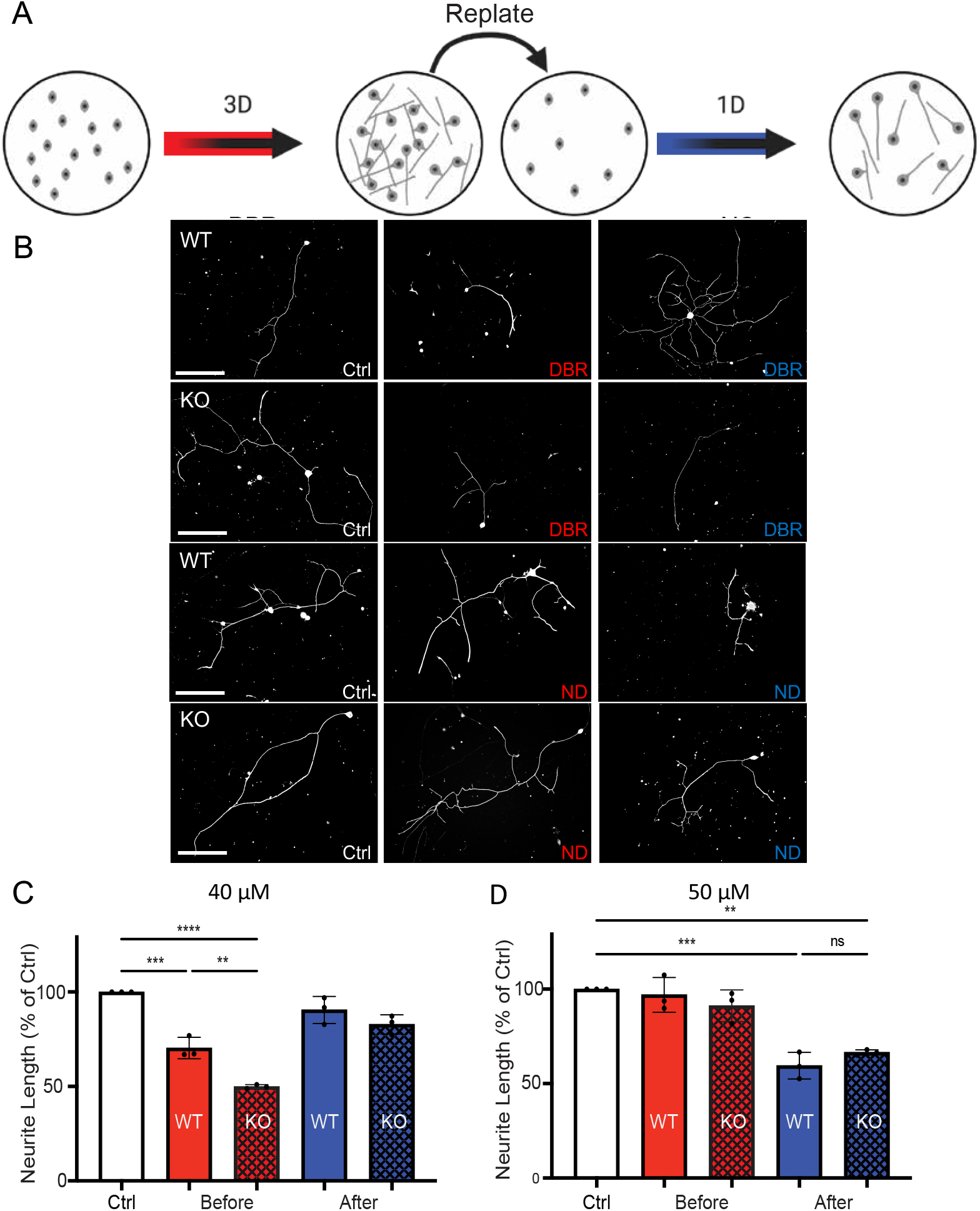
Caldendrin KO DRGNs exhibit stronger transcription-dependent neurite regeneration than WT DRGNs following *in vitro* axotomy. (A) Schematic of the experimental design for the same described in Fig.3. (B) Representative images of DRGNs from WT or KO male mice cultured with control medium (Ctrl) or medium containing 5,6-dichlorobenzimidazole riboside (DBR, 40 µM) or nocodazole (ND, 50 µM) before or after replating. Scale bars, 500 µm. (C,D) Longest neurite was measured as in Fig.3 and plotted for control and experimental groups. Each point represents the average obtained from n = 33 DRGNs. Columns and error bars represent mean ± SEM. **p<0.01, ***p<0.001, ****p<0.0001, determined by two-way ANOVA, Tukey’s multiple comparisons test.

GAP43 is the product of a RAG that regulates actin dynamics and facilitates neurite growth during neurodevelopment and regeneration (Chung et al., 2020). By western blot, we noted a time-dependent increase in GAP43 expression and corresponding decrease in caldendrin expression between 3 h and 3 d in WT cultures (Fig.5). Thus, downregulation of caldendrin may be necessary for induction of RAGs that support neurite growth.

**Figure 5.**
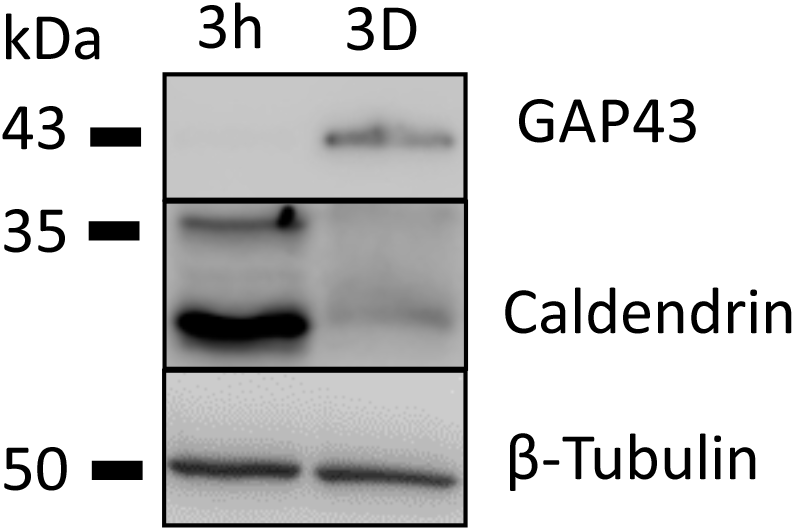
Caldendrin and GAP43 undergo opposing changes in expression with time in culture. DRGN cultures from WT male mice were lysed 3 h or 3 d after plating and subject to western blotting with antibodies against caldendrin, GAP43, and ß-Tubulin for loading control.

### Sex-dependent role of caldendrin in activity-dependent repression of neurite growth

Although it promotes neurite growth in some neurons (Goldberg et al., 2002; Singh & Miller, 2005), electrical activity serves as a brake on neurite growth in DRGNs (Fields et al., 1990). through activation of Ca_v_1 L-type Ca^2+^ channels (Enes et al., 2010; Robson & Burgoyne, 1989). Since caldendrin interacts with and potentiates the opening of Ca_v_1 channels (Tippens & Lee, 2007), caldendrin KO DRGNs are predicted to be less sensitive to the growth-inhibitory effects of electrical activity. To test this, we compared the impact of a depolarizing concentration of K^+^ (40 mM, K_40_) on neurite regeneration in WT and caldendrin KO DRGNs. As expected, K_40_ caused a dramatic reduction in the number of WT DRGNs with neurites at both 18 h and 24 h *in vitro*, and this was abrogated by co-treatment with the Ca_v_1 blockers isradipine and nimodipine. In contrast, there was no significant effect of K_40_ on DRGNs from caldendrin KO females at either time point (Fig.6A,B). Surprisingly, K_40_ strongly diminished the number of DRGNs with neurites in cultures from caldendrin KO as well as WT males (Fig.6C). To further probe these sex-specific actions of caldendrin, we compared the effects of K_40_ administered in the *in vitro* axotomy assay (Fig.7A). While K_40_ administered prior to replating strongly inhibited neurite regeneration in a manner that was reversed by nimodipine in cultures from WT females, this effect of K_40_ was absent in cultures from caldendrin KO females (Fig.7B). While K_40_ reduced neurite lengths when given before but not after replating in cultures from WT females, neurite lengths were not significantly different in cultures from caldendrin KO females regardless of K_40_ exposure (Fig.7C). Collectively, our findings suggest that caldendrin couples Ca_v_1 channels to activity-dependent transcriptional pathways that repress neurite regeneration in DRGNs in females but not in males.

**Figure 6.**
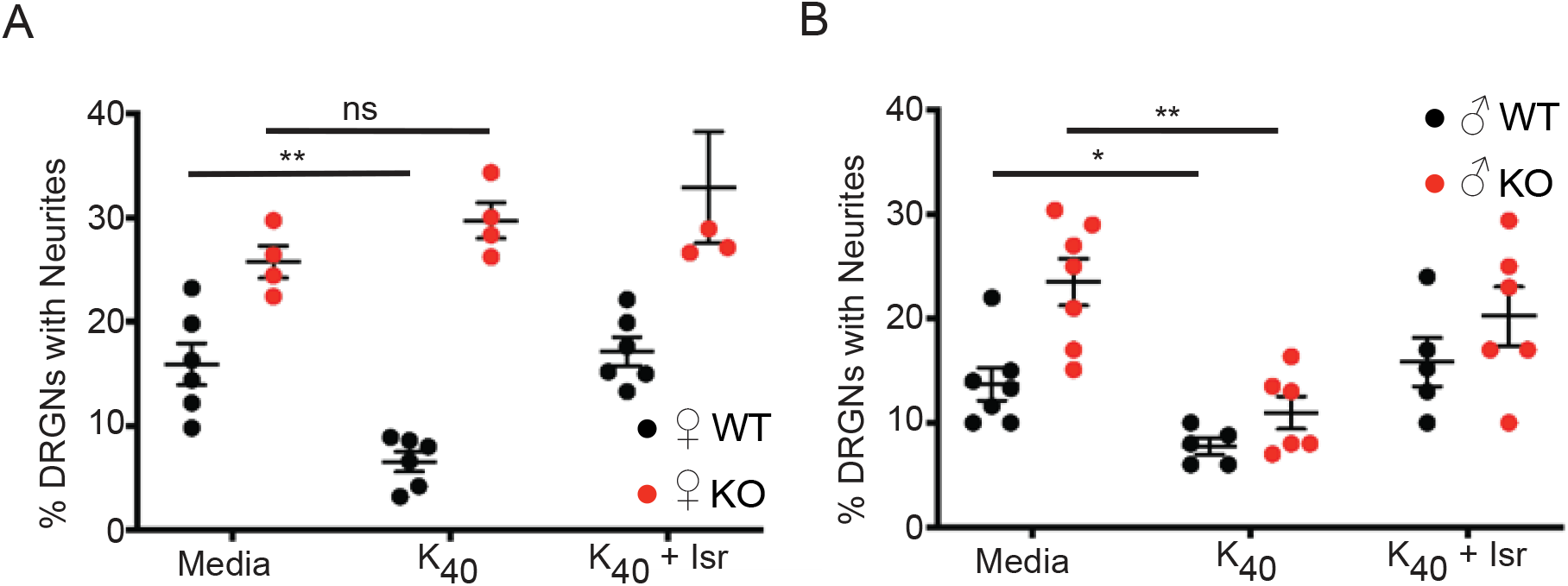
DRGNs from caldendrin KO females but not males are deficient in Ca_v_1-dependent neurite growth repression. (A,B) DRGNs from female (A) or male (B) mice were cultured for 18 h in the presence of control medium (Ctrl) or medium containing 40 mM KCl (K_40_) ± ISR (10 µM) prior to determination of % DRGNs with neurites as in Fig.2. Each point represents average determined from a total of ∼ 300 DRGNs per culture. Bars represent mean ±SEM. *p<0.05, **p<0.001, by one-way ANOVA with Dunnett’s multiple comparisons test.

**Figure 7.**
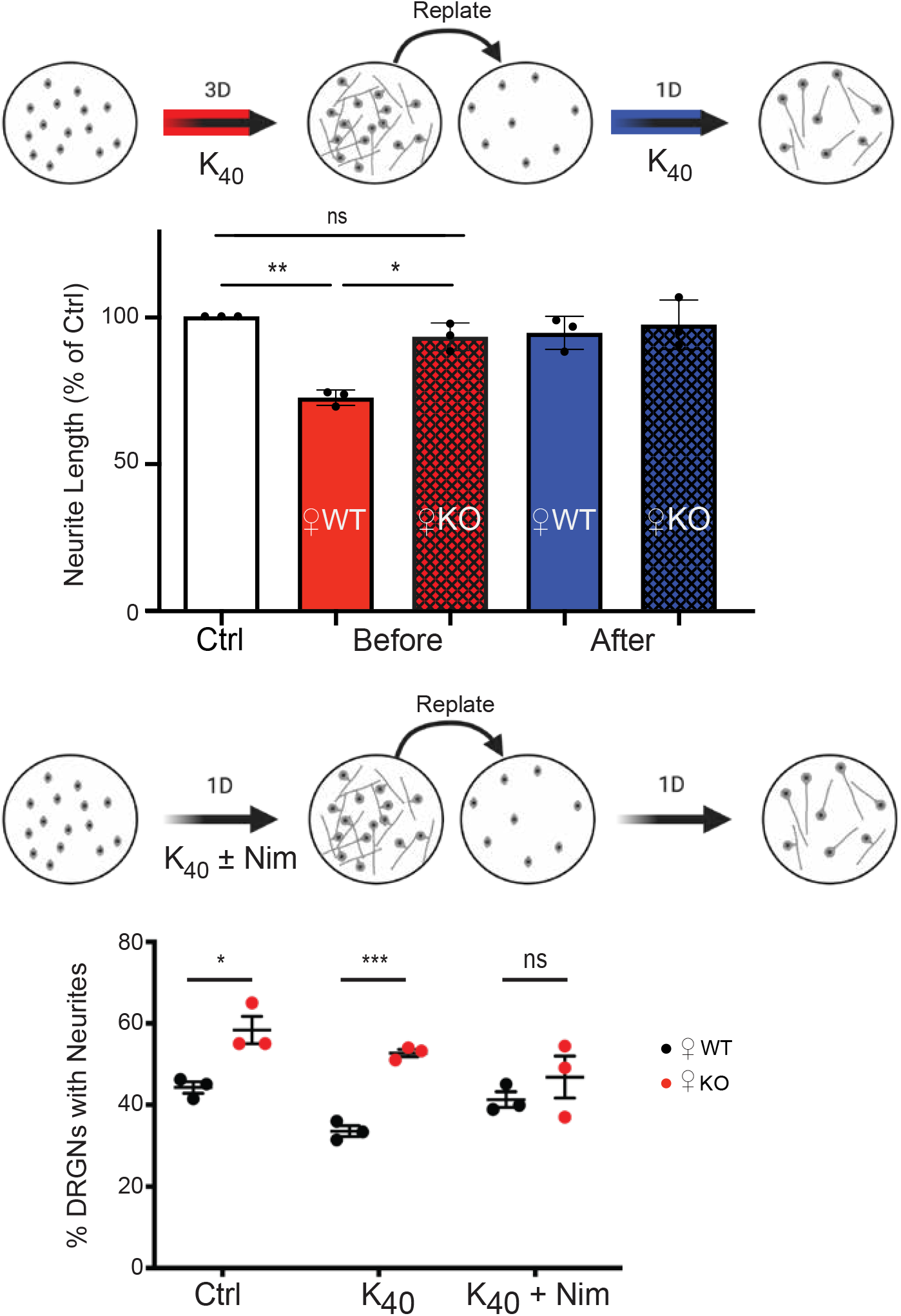
Coupling of Ca_v_1 channels to transcription-dependent repression of neurite regeneration following *in vitro* axotomy is disrupted in DRGNs from caldendrin KO females. (A) Schematic of the experimental design indicating exposure of cultures to K_40_ either before or after replating. (B) Longest neurite was measured as in Fig.3 and plotted for control and experimental groups in which WT or KO cultures were exposed to K_40_ before or after replating. *p=0.01, **p=0.002 by two-way ANOVA, Tukey’s multiple comparisons test. (C) Same as in A except that cultures were exposed to K_40_ ± nimodipine (Nim, 10 µM) before or after replating. (D) % DRGNs with neurites was determined as in Fig.2, each data point represents the average determined from a total of 400 DRGNs per culture. *p< 0.05, ***p= 0.0003, determined by t-test.

## DISCUSSION

During embryonic development, neurons in the central nervous system extend their axons often over long distances to innervate their targets but later lose this capability, presumably because regenerative signaling pathways become silenced (Yiera & Bradke, 2006). In DRGNs, severing the peripheral neurite can re-activate the genetic program that promotes growth of both the peripherally and centrally directed neurite (Smith & Skene, 1997). Thus, DRGNs represent an excellent model in which to study the mechanisms that normally suppress neurite regeneration. Upon dissociation from adult animals, these neurons readily regrow neurites within hours in culture. Our findings that, compared to DRGNs from WT mice, those from caldendrin KO mice had more robustly regenerate their neurites in standard cultures (Fig.2) and following *in vitro* axotomy (Fig.3) implicate caldendrin in the pathway that opposes the regenerative neurite growth in these primary sensory neurons.

Like CaM, caldendrin can bind to and regulate a variety of effector molecules that regulate Ca^2+^ signaling pathways that impinge on neurite growth dynamics. A major candidate in this regard is the Ca_v_1 channel, which mediates the inhibitory effects of electrical activity on neurite growth (Enes et al., 2010; Robson & Burgoyne, 1989). Caldendrin and shorter CaBP1 variants bind to the pore-forming subunit of the major Ca_v_1 subtypes expressed in the nervous system, Ca_v_1.2 and Ca_v_1.3, and this interaction potentiates channel opening (Cui et al., 2007; Tippens & Lee, 2007; Zhou et al., 2004; Zhou et al., 2005). Thus, Ca_v_1-dependent Ca^2+^ signals are expected to be abnormally low in the absence of caldendrin, which we have previously shown to be the case in cochlear spiral ganglion neurons of neonatal caldendrin KO mice (T. Yang et al., 2018). If Ca_v_1 were similarly depressed in caldendrin KO DRGNs, Ca_v_1 Ca^2+^ signals may be insufficient to oppose the transcription of RAGs that such as GAP43. In this regard, it is noteworthy that caldendrin levels were low in WT DRGNs at timepoints when GAP43 was high (Fig.5). Our findings show DBR had a stronger effect in repressing neurite growth in caldendrin KO than in WT cultures (Fig.4) supports the existence of heightened pro-regenerative transcriptional activity in caldendrin KO DRGNs. We speculate that caldendrin controls the activity/expression of transcriptional repressors such as RE1 Silencing Transcription Factor (REST), which suppress transcription of genes that promote axon outgrowth (Oh et al., 2018). In hippocampal neurons, caldendrin binds to and regulates the nuclear localization of Jacob, a protein that controls cAMP-response element binding protein (CREB)-dependent synaptic remodeling (Dieterich et al., 2008). Interactions between CREB and REST pathways have been reported in neuronal gene regulation networks (Wu & Xie, 2006) but whether these interactions occur in DRGNs has not been investigated.

In DRGN cultures, the involvement of Ca_v_1 Ca^2+^ signals in neurite growth is readily revealed by depolarizing concentrations of K^+^ which repress neurite growth in a manner that is reversed by Ca_v_1 blockers (Enes et al., 2010; Robson & Burgoyne, 1989). The absence of any effect of K_40_ on neurite growth in caldendrin KO DRGNs in conventional culture (Fig.6A) and following *in vitro* axotomy (Fig.7) indicates a critical role for caldendrin in coupling Ca_v_1 channels to activity-dependent neurite growth repression (Fig.6A). The fact that this was only applicable to cultures from female and not male caldendrin KO mice (Fig.6B) was intriguing considering that sex differences have been noted in the transcriptome of mouse DRGNs (Mecklenburg et al., 2020) and in the responses to peripheral nerve injury in animals and humans (Chernov et al., 2020; Franco-Enzastiga et al., 2021; Juarez et al., 2019). This result was not due to major differences in caldendrin protein levels in WT males and females (Fig.1B). One possibility is that circulating sex steroids in female mice such as estradiol somehow promote caldendrin interactions with Ca_v_1 channels. Estradiol has cell type-specific modulatory effects on Ca_v_1 channels (Lai et al., 2019; Sarkar et al., 2008; Vega-Vela et al., 2017; X. Yang et al., 2018) and so could modify channel gating mechanisms that are targeted by caldendrin.

An important follow-up question is how does caldendrin regulate neurite growth in DRGNs from males if Ca_v_1 channels are not involved? Like the shorter CaBP1 variants, caldendrin could bind to and inhibit inositol 1,4,5 trisphosphate receptors (IP_3_Rs) (Haynes et al., 2004; Kasri et al., 2004) and therefore IP_3_R-mediated Ca^2+^ signals that are known to enhance DRGN neurite outgrowth (Takei et al., 1998). Alternatively, caldendrin might interact with the actin-binding protein cortactin (Mikhaylova et al., 2018) and A-kinase anchoring proteins (AKAPs) (Gorny et al., 2012), both of which have been implicated in neurite growth regulation (Boczek et al., 2021; Kubo et al., 2015).

In summary, we have established a novel role for caldendrin in a sexually dimorphic signaling pathway that represses neurite regeneration in DRGNs *in vitro*. Establishing the significance of this pathway in peripheral nerve injury models in vivo, as well as an understanding of the mechanisms differentiating this pathway in males and females, remain important challenges for future studies.

## Abbreviations

CaBP1: calcium binding protein 1
DRGN: dorsal root ganglion neuron
KO: knock out
NF200: neurofilament 200
PBS: phosphate buffered saline
PFA: paraformaldehyde
PNS: peripheral nervous system
RAG: regeneration associated gene
WT: wild type.

## Notes

### Competing Interest Statement

The authors have declared no competing interest.

## REFERENCES

Boczek, T., Yu, Q., Zhu, Y., Dodge-Kafka, K. L., Goldberg, J. L., & Kapiloff, M. S. (2021). cAMP at Perinuclear mAKAPalpha Signalosomes Is Regulated by Local Ca(2+) Signaling in Primary Hippocampal Neurons. eNeuro, 8(1). https://doi.org/10.1523/ENEURO.0298-20.2021

Chernov, A. V., Hullugundi, S. K., Eddinger, K. A., Dolkas, J., Remacle, A. G., Angert, M., James, B. P., Yaksh, T. L., Strongin, A. Y., & Shubayev, V. I. (2020). A myelin basic protein fragment induces sexually dimorphic transcriptome signatures of neuropathic pain in mice. Journal of Biological Chemistry, 295(31), 10807–10821. https://doi.org/10.1074/jbc.RA120.013696

Chung, D., Shum, A., & Caraveo, G. (2020). GAP-43 and BASP1 in Axon Regeneration: Implications for the Treatment of Neurodegenerative Diseases. Front Cell Dev Biol, 8, 567537. https://doi.org/10.3389/fcell.2020.567537

Costigan, M., Befort, K., Karchewski, L., Griffin, R. S., D’Urso, D., Allchorne, A., Sitarski, J., Mannion, J. W., Pratt, R. E., & Woolf, C. J. (2002). Replicate high-density rat genome oligonucleotide microarrays reveal hundreds of regulated genes in the dorsal root ganglion after peripheral nerve injury. BMC Neurosci, 3, 16. https://doi.org/10.1186/1471-2202-3-16

Cui, G., Meyer, A. C., Calin-Jageman, I., Neef, J., Haeseleer, F., Moser, T., & Lee, A. (2007). Ca^2+^-binding proteins tune Ca^2+^-feedback to Cav1.3 channels in auditory hair cells. The Journal of physiology, 585, 791-803. 17947313

Dieterich, D. C., Karpova, A., Mikhaylova, M., Zdobnova, I., Konig, I., Landwehr, M., Kreutz, M., Smalla, K. H., Richter, K., Landgraf, P., Reissner, C., Boeckers, T. M., Zuschratter, W., Spilker, C., Seidenbecher, C. I., Garner, C. C., Gundelfinger, E. D., & Kreutz, M. R. (2008). Caldendrin-Jacob: a protein liaison that couples NMDA receptor signalling to the nucleus. PLoS Biol, 6(2), e34. https://doi.org/10.1371/journal.pbio.0060034

Enes, J., Langwieser, N., Ruschel, J., Carballosa-Gonzalez, M. M., Klug, A., Traut, M. H., Ylera, B., Tahirovic, S., Hofmann, F., Stein, V., Moosmang, S., Hentall, I. D., & Bradke, F. (2010). Electrical activity suppresses axon growth through Ca(v)1.2 channels in adult primary sensory neurons [Research Support, Non-U.S. Gov’t]. Current biology : CB, 20(13), 1154–1164. https://doi.org/10.1016/j.cub.2010.05.055

Fields, R. D., Neale, E. A., & Nelson, P. G. (1990). Effects of patterned electrical activity on neurite outgrowth from mouse sensory neurons. The Journal of Neuroscience, 10(9), 2950–2964. https://www.ncbi.nlm.nih.gov/pubmed/2398369

Franco-Enzastiga, U., Garcia, G., Murbartian, J., Gonzalez-Barrios, R., Salinas-Abarca, A. B., Sanchez-Hernandez, B., Tavares-Ferreira, D., Herrera, L. A., Barragan-Iglesias, P., Delgado-Lezama, R., Price, T. J., & Granados-Soto, V. (2021). Sex-dependent pronociceptive role of spinal alpha5 -GABAA receptor and its epigenetic regulation in neuropathic rodents. J Neurochem, 156(6), 897–916. https://doi.org/10.1111/jnc.15140

Goldberg, J. L., Espinosa, J. S., Xu, Y., Davidson, N., Kovacs, G. T., & Barres, B. A. (2002). Retinal ganglion cells do not extend axons by default: promotion by neurotrophic signaling and electrical activity. Neuron, 33(5), 689–702. https://www.ncbi.nlm.nih.gov/pubmed/11879647

Gomez, T. M., & Zheng, J. Q. (2006). The molecular basis for calcium-dependent axon pathfinding. Nature reviews. Neuroscience, 7(2), 115–125. https://doi.org/10.1038/nrn1844

Gorny, X., Mikhaylova, M., Seeger, C., Reddy, P. P., Reissner, C., Schott, B. H., Helena Danielson, U., Kreutz, M. R., & Seidenbecher, C. (2012). AKAP79/150 interacts with the neuronal calcium-binding protein caldendrin [In Vitro Research Support, Non-U.S. Gov’t]. J Neurochem, 122(4), 714–726. https://doi.org/10.1111/j.1471-4159.2012.07828.x

Haeseleer, F., Imanishi, Y., Maeda, T., Possin, D. E., Maeda, A., Lee, A., Rieke, F., & Palczewski, K. (2004). Essential role of Ca^2+^-binding protein 4, a Cav1.4 channel regulator, in photoreceptor synaptic function. Nature Neuroscience, 7(10), 1079-1087. 15452577

Haeseleer, F., Sokal, I., Verlinde, C. L., Erdjument-Bromage, H., Tempst, P., Pronin, A. N., Benovic, J. L., Fariss, R. N., & Palczewski, K. (2000). Five members of a novel Ca^2+^-binding protein (CABP) subfamily with similarity to calmodulin. Journal of Biological Chemistry, 275(2), 1247-1260. 10625670

Haynes, L. P., Tepikin, A. V., & Burgoyne, R. D. (2004). Calcium-binding protein 1 is an inhibitor of agonist-evoked, inositol 1,4,5-trisphosphate-mediated calcium signaling. Journal of Biological Chemistry, 279(1), 547-555. 14570872

Henley, J., & Poo, M. M. (2004). Guiding neuronal growth cones using Ca^2+^ signals. Trends Cell Biol, 14(6), 320–330. https://doi.org/10.1016/j.tcb.2004.04.006

Huebner, E. A., Budel, S., Jiang, Z., Omura, T., Ho, T. S., Barrett, L., Merkel, J. S., Pereira, L. M., Andrews, N. A., Wang, X., Singh, B., Kapur, K., Costigan, M., Strittmatter, S. M., & Woolf, C. J. (2019). Diltiazem Promotes Regenerative Axon Growth. Mol Neurobiol, 56(6), 3948–3957. https://doi.org/10.1007/s12035-018-1349-5

Huebner, E. A., & Strittmatter, S. M. (2009). Axon regeneration in the peripheral and central nervous systems. Results Probl Cell Differ, 48, 339–351. https://doi.org/10.1007/400_2009_19

Juarez, I., Morales-Medina, J. C., Flores-Tochihuitl, J., Juarez, G. S., Flores, G., & Oseki, H. C. (2019). Tooth pulp injury induces sex-dependent neuronal reshaping in the ventral posterolateral nucleus of the rat thalamus. J Chem Neuroanat, 96, 16–21. https://doi.org/10.1016/j.jchemneu.2018.10.007

Kasri, N. N., Holmes, A. M., Bultynck, G., Parys, J. B., Bootman, M. D., Rietdorf, K., Missiaen, L., McDonald, F., De Smedt, H., Conway, S. J., Holmes, A. B., Berridge, M. J., & Roderick, H. L. (2004). Regulation of InsP3 receptor activity by neuronal Ca^2+^-binding proteins. The EMBO journal, 23(2), 312-321. 14685260

Kim, K. Y., Scholl, E. S., Liu, X., Shepherd, A., Haeseleer, F., & Lee, A. (2014). Localization and expression of CaBP1/caldendrin in the mouse brain. Neuroscience, 268, 33–47. https://doi.org/10.1016/j.neuroscience.2014.02.052

Kubo, Y., Baba, K., Toriyama, M., Minegishi, T., Sugiura, T., Kozawa, S., Ikeda, K., & Inagaki, N. (2015). Shootin1-cortactin interaction mediates signal-force transduction for axon outgrowth. The Journal of cell biology, 210(4), 663–676. https://doi.org/10.1083/jcb.201505011

Lai, Y. J., Zhu, B. L., Sun, F., Luo, D., Ma, Y. L., Luo, B., Tang, J., Xiong, M. J., Liu, L., Long, Y., Hu, X. T., He, L., Deng, X. J., Zhang, J. H., Yang, J., Yan, Z., & Chen, G. J. (2019). Estrogen receptor alpha promotes Cav1.2 ubiquitination and degradation in neuronal cells and in APP/PS1 mice. Aging Cell, 18(4), e12961. https://doi.org/10.1111/acel.12961

Laube, G., Seidenbecher, C. I., Richter, K., Dieterich, D. C., Hoffmann, B., Landwehr, M., Smalla, K. H., Winter, C., Bockers, T. M., Wolf, G., Gundelfinger, E. D., & Kreutz, M. R. (2002). The neuron-specific Ca^2+^-binding protein caldendrin: gene structure, splice isoforms, and expression in the rat central nervous system. Mol. Cell. Neurosci., 19(3), 459-475. 11906216

Lawson, S. N., & Waddell, P. J. (1991). Soma neurofilament immunoreactivity is related to cell size and fibre conduction velocity in rat primary sensory neurons. J Physiol, 435, 41–63. https://doi.org/10.1113/jphysiol.1991.sp018497

Lin, Y. T., & Chen, J. C. (2018). Dorsal Root Ganglia Isolation and Primary Culture to Study Neurotransmitter Release. Journal of visualized experiments : JoVE(140). https://doi.org/10.3791/57569

Mecklenburg, J., Zou, Y., Wangzhou, A., Garcia, D., Lai, Z., Tumanov, A. V., Dussor, G., Price, T. J., & Akopian, A. N. (2020). Transcriptomic sex differences in sensory neuronal populations of mice. Scientific reports, 10(1), 15278. https://doi.org/10.1038/s41598-020-72285-z

Mikhaylova, M., Bar, J., van Bommel, B., Schatzle, P., YuanXiang, P., Raman, R., Hradsky, J., Konietzny, A., Loktionov, E. Y., Reddy, P. P., Lopez-Rojas, J., Spilker, C., Kobler, O., Raza, S. A., Stork, O., Hoogenraad, C. C., & Kreutz, M. R. (2018). Caldendrin Directly Couples Postsynaptic Calcium Signals to Actin Remodeling in Dendritic Spines. Neuron, 97(5), 1110–1125 e1114. https://doi.org/10.1016/j.neuron.2018.01.046

Moskowitz, P. F., Smith, R., Pickett, J., Frankfurter, A., & Oblinger, M. M. (1993). Expression of the class III beta-tubulin gene during axonal regeneration of rat dorsal root ganglion neurons. Journal of neuroscience research, 34(1), 129–134. https://doi.org/10.1002/jnr.490340113

Oh, Y. M., Mahar, M., Ewan, E. E., Leahy, K. M., Zhao, G., & Cavalli, V. (2018). Epigenetic regulator UHRF1 inactivates REST and growth suppressor gene expression via DNA methylation to promote axon regeneration. Proc Natl Acad Sci U S A, 115(52), E12417–E12426. https://doi.org/10.1073/pnas.1812518115

Popko, J., Fernandes, A., Brites, D., & Lanier, L. M. (2009). Automated analysis of NeuronJ tracing data. Cytometry A, 75(4), 371–376. https://doi.org/10.1002/cyto.a.20660

Robson, S. J., & Burgoyne, R. D. (1989). L-type calcium channels in the regulation of neurite outgrowth from rat dorsal root ganglion neurons in culture. Neurosci Lett, 104(1-2), 110–114. https://www.ncbi.nlm.nih.gov/pubmed/2554216

Saijilafu Hur, E.M., Liu, C. M., Jiao, Z., Xu, W. L., & Zhou, F. Q. (2013). PI3K-GSK3 signalling regulates mammalian axon regeneration by inducing the expression of Smad1. Nat Commun, 4, 2690. https://doi.org/10.1038/ncomms3690

Sarkar, S. N., Huang, R. Q., Logan, S. M., Yi, K. D., Dillon, G. H., & Simpkins, J. W. (2008). Estrogens directly potentiate neuronal L-type Ca^2+^ channels. Proc Natl Acad Sci U S A, 105(39), 15148–15153. https://doi.org/10.1073/pnas.0802379105

Scheib, J., & Hoke, A. (2013). Advances in peripheral nerve regeneration. Nat Rev Neurol, 9(12), 668–676. https://doi.org/10.1038/nrneurol.2013.227

Schreyer, D. J., & Skene, J. H. (1991). Fate of GAP-43 in ascending spinal axons of DRG neurons after peripheral nerve injury: delayed accumulation and correlation with regenerative potential. The Journal of Neuroscience, 11(12), 3738–3751. https://www.ncbi.nlm.nih.gov/pubmed/1836017

Seidenbecher, C. I., Langnaese, K., Sanmarti-Vila, L., Boeckers, T. M., Smalla, K. H., Sabel, B. A., Garner, C. C., Gundelfinger, E. D., & Kreutz, M. R. (1998). Caldendrin, a novel neuronal calcium-binding protein confined to the somato-dendritic compartment. Journal of Biological Chemistry, 273(33), 21324-21331. 9694893

Singh, K. K., & Miller, F. D. (2005). Activity regulates positive and negative neurotrophin-derived signals to determine axon competition. Neuron, 45(6), 837–845. https://doi.org/10.1016/j.neuron.2005.01.049

Smith, D. S., & Skene, J. H. (1997). A transcription-dependent switch controls competence of adult neurons for distinct modes of axon growth. J Neurosci, 17(2), 646–658. https://www.ncbi.nlm.nih.gov/pubmed/8987787

Takei, K., Shin, R. M., Inoue, T., Kato, K., & Mikoshiba, K. (1998). Regulation of nerve growth mediated by inositol 1,4,5-trisphosphate receptors in growth cones. Science, 282(5394), 1705–1708. https://doi.org/10.1126/science.282.5394.1705

Tippens, A. L., & Lee, A. (2007). Caldendrin: a neuron-specific modulator of Cav1.2 (L-type) Ca^2+^ channels. Journal of Biological Chemistry, 282, 8464-8473. 17224447

Vega-Vela, N. E., Osorio, D., Avila-Rodriguez, M., Gonzalez, J., Garcia-Segura, L. M., Echeverria, V., & Barreto, G. E. (2017). L-Type Calcium Channels Modulation by Estradiol. Mol Neurobiol, 54(7), 4996–5007. https://doi.org/10.1007/s12035-016-0045-6

Wu, H., Petitpre, C., Fontanet, P., Sharma, A., Bellardita, C., Quadros, R. M., Jannig, P. R., Wang, Y., Heimel, J. A., Cheung, K. K. Y., Wanderoy, S., Xuan, Y., Meletis, K., Ruas, J., Gurumurthy, C. B., Kiehn, O., Hadjab, S., & Lallemend, F. (2021). Distinct subtypes of proprioceptive dorsal root ganglion neurons regulate adaptive proprioception in mice. Nature communications, 12(1), 1026. https://doi.org/10.1038/s41467-021-21173-9

Wu, J., & Xie, X. (2006). Comparative sequence analysis reveals an intricate network among REST, CREB and miRNA in mediating neuronal gene expression. Genome Biol, 7(9), R85. https://doi.org/10.1186/gb-2006-7 9-r85

Yang, T., Choi, J. E., Soh, D., Tobin, K., Joiner, M. L., Hansen, M., & Lee, A. (2018). CaBP1 regulates Cav1 L-type Ca(2+) channels and their coupling to neurite growth and gene transcription in mouse spiral ganglion neurons. Mol Cell Neurosci, 88, 342–352. https://doi.org/10.1016/j.mcn.2018.03.005

Yang, T., Scholl, E. S., Pan, N., Fritzsch, B., Haeseleer, F., & Lee, A. (2016). Expression and Localization of CaBP Ca^2+^ Binding Proteins in the Mouse Cochlea. PLoS One, 11(1), e0147495. https://doi.org/10.1371/journal.pone.0147495

Yang, X., Mao, X., Xu, G., Xing, S., Chattopadhyay, A., Jin, S., & Salama, G. (2018). Estradiol up-regulates L-type Ca(2+) channels via membrane-bound estrogen receptor/phosphoinositide-3-kinase/Akt/cAMP response element-binding protein signaling pathway. Heart Rhythm, 15(5), 741–749. https://doi.org/10.1016/j.hrthm.2018.01.019

Yiera, B., & Bradke, F. (2006). Stimulating Intrinsic Growth Potential in Mammalian Neurons. In C.G. Becker & T. Becker (Eds.), Model Organisms in Spinal Cord Regeneration Wiley-VCH Verlag GmbH & Co. KGaA.

Zheng, Y., Liu, P., Bai, L., Trimmer, J. S., Bean, B. P., & Ginty, D. D. (2019). Deep Sequencing of Somatosensory Neurons Reveals Molecular Determinants of Intrinsic Physiological Properties. Neuron, 103(4), 598–616 e597. https://doi.org/10.1016/j.neuron.2019.05.039

Zhou, H., Kim, S. A., Kirk, E. A., Tippens, A. L., Sun, H., Haeseleer, F., & Lee, A. (2004). Ca^2+^-binding protein-1 facilitates and forms a postsynaptic complex with Cav1.2 (L-type) Ca^2+^ channels. The Journal of Neuroscience, 24(19), 4698-4708. 15140941

Zhou, H., Yu, K., McCoy, K. L., & Lee, A. (2005). Molecular mechanism for divergent regulation of Cav1.2 Ca^2+^ channels by calmodulin and Ca^2+^-binding protein-1. Journal of Biological Chemistry, 280(33), 29612-29619. 15980432

